# Clearance of *Clostridioides difficile* colonization is associated with antibiotic-specific bacterial changes

**DOI:** 10.1101/2020.11.16.386086

**Authors:** Nicholas A. Lesniak, Alyxandria M. Schubert, Hamide Sinani, Patrick D. Schloss

## Abstract

The gut bacterial community prevents many pathogens from colonizing the intestine. Previous studies have associated specific bacteria with clearing *Clostridioides difficile* colonization across different community perturbations. However, those bacteria alone have been unable to clear *C. difficile* colonization. To elucidate the changes necessary to clear colonization, we compared differences in bacterial abundance between communities able and unable to clear *C. difficile* colonization. We treated mice with titrated doses of antibiotics prior to *C. difficile* challenge which resulted in no colonization, colonization and clearance, or persistent colonization. Previously, we observed that clindamycin-treated mice were susceptible to colonization but spontaneously cleared *C. difficile.* Therefore, we investigated whether other antibiotics would show the same result. We found reduced doses of cefoperazone and streptomycin permitted colonization and clearance of *C. difficile.* Mice that cleared colonization had antibiotic-specific community changes and predicted interactions with *C. difficile.* Clindamycin treatment led to a bloom in populations related to *Enterobacteriaceae.* Clearance of *C. difficile* was concurrent with the reduction of those blooming populations and the restoration of community members related to the *Porphyromonadaceae* and *Bacteroides.* Cefoperazone created a susceptible community characterized by a drastic reduction in the community diversity, interactions, and a sustained increase in abundance of many facultative anaerobes. Lastly, clearance in streptomycin-treated mice was associated with the recovery of multiple members of the *Porphyromonadaceae*, with little overlap in the specific *Porphyromonadaceae* observed in the clindamycin treatment. Further elucidation of how *C. difficile* colonization is cleared from different gut bacterial communities will improve *C. difficile* infection treatments.

**Importance:** The community of microorganisms, known as the microbiota, in our intestines prevents pathogens, such as *C. difficile*, from establishing themselves and causing infection. This is known as colonization resistance. However, when a person takes antibiotics, their gut microbiota is disturbed. This disruption allows *C. difficile* to colonize. *C. difficile* infections (CDI) are primarily treated with antibiotics, which frequently leads to recurrent infections because the microbiota have not yet returned to a resistant state. The infection cycle often ends when the fecal microbiota from a presumed resistant person are transplanted into the susceptible person. Although this treatment is highly effective, we do not understand the mechanism of resistance. We hope to improve the treatment of CDI through elucidating how the bacterial community eliminates *C. difficile* colonization. We found *C. difficile* was able to colonize susceptible mice but was spontaneously eliminated in an antibiotic-treatment specific manner. These data indicate each community had different requirements for clearing colonization. Understanding how different communities clear colonization will reveal targets to improve CDI treatments.

## Introduction

A complex consortium of bacteria and microbes that inhabits our gut, known as the microbiota, prevent pathogens from colonizing and causing disease. This protection, known as colonization resistance, is mediated through many mechanisms such as activating host immune responses, competing for nutrients, producing antimicrobials, and contributing to the maintenance of the mucosal barrier (1). However, perturbations to the intestinal community or these functions opens the possibility that a pathogen can colonize (2). For example, the use of antibiotics perturb the gut microbiota and can lead to *Clostridioides difficile* infection (CDI).

CDI is especially problematic due to its burden on the healthcare system (3, 4). *C. difficile* can cause severe disease, such as toxic megacolon, diarrhea, and death (5). CDI is primarily treated with antibiotics (6). CDIs recalcitrant to antibiotics are eliminated by restoring the community with a fecal microbiota transplant (FMT), returning the perturbed community to a healthier protective state (7, 8). However, FMTs are not always effective against CDI and have the risk of transferring a secondary infection (9, 10). Therefore, we need to better understand how the microbiota clears the infection to develop more effective treatments.

Previous research has shown that the microbiota affects *C. difficile* colonization. Mouse models have identified potential mechanisms of colonization resistance such as bile salt metabolism and nutrient competition (11–14). However, studies that have restored those functions were unable to restore complete resistance (15, 16). This could be attributed to the complexity of the community and the mechanisms of colonization resistance (17, 18). We previously showed that when *C. difficile* colonizes different antibiotic-treated murine communities it modifies its metabolism to fit each specific environment (14, 19, 20). Therefore, we have investigated the bacterial community dynamics concurrent with *C. difficile* elimination across uniquely perturbed communities.

Jenior et al. (20) observed that clindamycin-treated mice spontaneously cleared *C. difficile* colonization whereas mice treated with cefoperazone and streptomycin did not. Here, we continued to explore the different effects these three antibiotics have on *C. difficile* colonization. The purpose of this study was to elucidate the gut bacterial community changes concurrent with elimination of *C. difficile* colonization. We hypothesized that each colonized community has perturbation-specific susceptibilities and requires specific changes to clear the pathogen. To induce a less severe perturbation, we reduced the doses of cefoperazone and streptomycin. This resulted in communities that were initially colonized to a high level (>10^6^ CFU/g feces) and then spontaneously cleared *C. difficile.* We found each antibiotic resulted in unique changes in the microbiota that were associated with the persistence or clearance of *C. difficile.* These data further support the hypothesis that *C. difficile* can exploit numerous niches in perturbed communities.

## Results

### Reduced doses of cefoperazone and streptomycin allowed communities to spontaneously clear *C. difficile* colonization

To understand the dynamics of colonization and clearance of *C. difficile*, we first identified conditions which would allow colonization and clearance. Beginning with clindamycin, mice were treated with an intraperitoneal injection of clindamycin (10 mg/kg) one day prior to challenge with *C. difficile.* All mice (N = 11) were colonized to a high level (median CFU = 3.07×10^7^) the next day and cleared the colonization within 10 days; 6 mice cleared *C. difficile* within 6 days (Figure 1A). Previous *C. difficile* infection models using cefoperazone and streptomycin have not demonstrated clearance. So we next explored whether cefoperazone and streptomycin could permit colonization and subsequent clearance with lower doses. We began with replicating the previously established *C. difficile* infection models using these antibiotics (20). We treated mice with cefoperazone or streptomycin in their drinking water for 5 days (0.5 mg/mL and 5 mg/mL, respectively) and then challenged them with *C. difficile*. For both antibiotics, *C. difficile* colonization was maintained for the duration of the experiment as previously demonstrated (Figure 1B-C) (20). Then we repeated the *C. difficile* challenge with reduced doses of the antibiotics (cefoperazone - 0.3 and 0.1 mg/mL; streptomycin - 0.5 and 0.1 mg/mL). For both antibiotic treatments, the lowest dose resulted in either no colonization (N = 8) or a transient, low level colonization (N = 8, median length = 1 day, median CFU/g =2.8×10^3^) (Figure 1B-C). The intermediate dose of both antibiotics resulted in a high level colonization (median CFU/g =3.5×10^6^) and half (N = 8 of 16) of the mice clearing the colonization within 10 days. Based on our previous research, which showed each of these antibiotics uniquely changed the microbiota, we hypothesized that the microbiota varied across these antibiotic treatments that resulted in colonization clearance.

**Figure 1.**
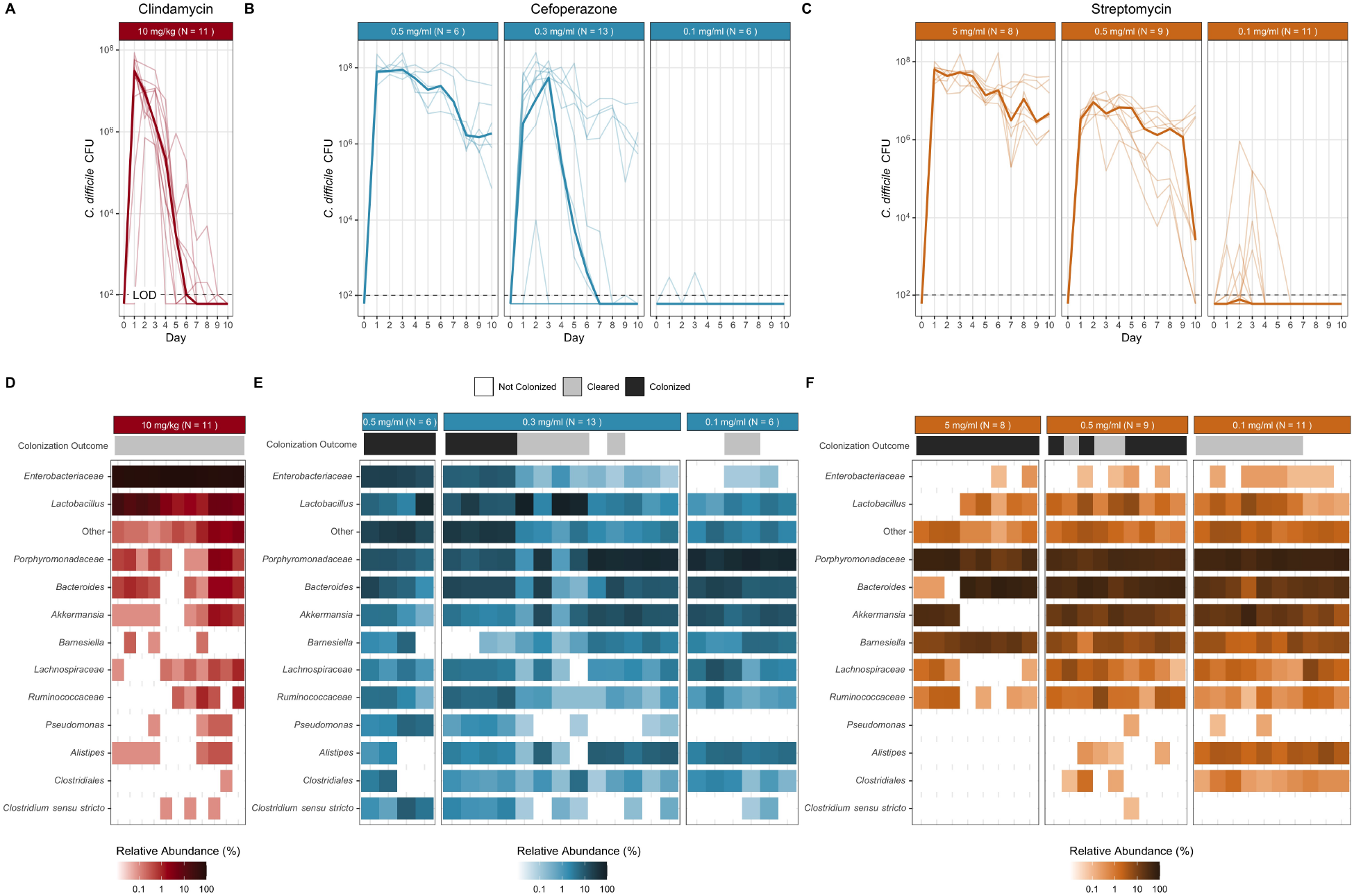
Reduced antibiotic doses permitted murine communities to be colonized and spontaneously clear that *C. difficile* colonization. (A-C) Daily CFU of *C. difficile* in fecal samples of mice treated with clindamycin, cefoperazone, or streptomycin from time of challenge (Day 0) through 10 days post infection (dpi). The bold line is the median CFU of the group and the transparent lines are the individual mice. (D-F) Relative abundance of twelve most abundant genera at the time of *C. difficile* challenge, all other genera grouped into Other. Each column is an individual mouse. LOD = Limit of detection. (clindamycin −10 mg/kg N =11; cefoperazone - 0.5 mg/mL N = 5, 0.3 mg/mL N = 9, 0.1 mg/mL N = 2; streptomycin - 5.0 mg/mL N = 8, 0.5 mg/mL N = 7, 0.1 mg/mL N = 7).

### Clearance of *C. difficile* was associated with antibiotic-specific changes to the microbiota

Beginning with the clindamycin-treated mice, we analyzed their fecal 16S rRNA gene sequences to identify the community features related to *C. difficile* colonization and clearance. First, we compared the most abundant bacterial genera of the communities at the time of *C. difficile* challenge. The clindamycin-treated mice became dominated by relatives of *Enterobacteriaceae* with a concurrent reduction in the other abundant genera, except for populations of *Lactobacillus* (Figure 1D, S1). These community changes permitted *C. difficile* to colonize all of these mice, but all of the mice were are also able to clear the colonization. We next investigated how the microbiota diversity related to *C. difficile* clearance. Clindamycin treatment decreased the alpha diversity (*P* < 0.05) and similarity to the pre-clindamycin community at the time of *C. difficile* challenge (P < 0.05) (Figure 2A). But it was not necessary to restore the community similarity to its initial state to clear *C. difficile*. Therefore we investigated the temporal differences in the abundance of the operational taxonomic units (OTUs) between the initial untreated community and post-clindamycin treatment at the time of challenge and between the time of challenge and the end of the experiment. Clindamycin treatment resulted in large decreases in 21 OTUs and a bloom of relatives of *Enterobacteriaceae* (Figure 4A). With the elimination of *C. difficile*, we observed a drastic reduction of the relatives of *Enterobacteriaceae* and recovery of 10 populations related to *Porphyromonadaceae, Bacteroides, Akkermansia, Lactobacillus, Bifidobacterium, Lachnospiraceae*, and *Clostridiales* (Figure 4A). Thus, clindamycin reduced most of the natural community allowing *C. difficile* to colonize. The recovery of only a small portion of the community was associated with eliminating the *C.difficile* population.

**Figure 2.**
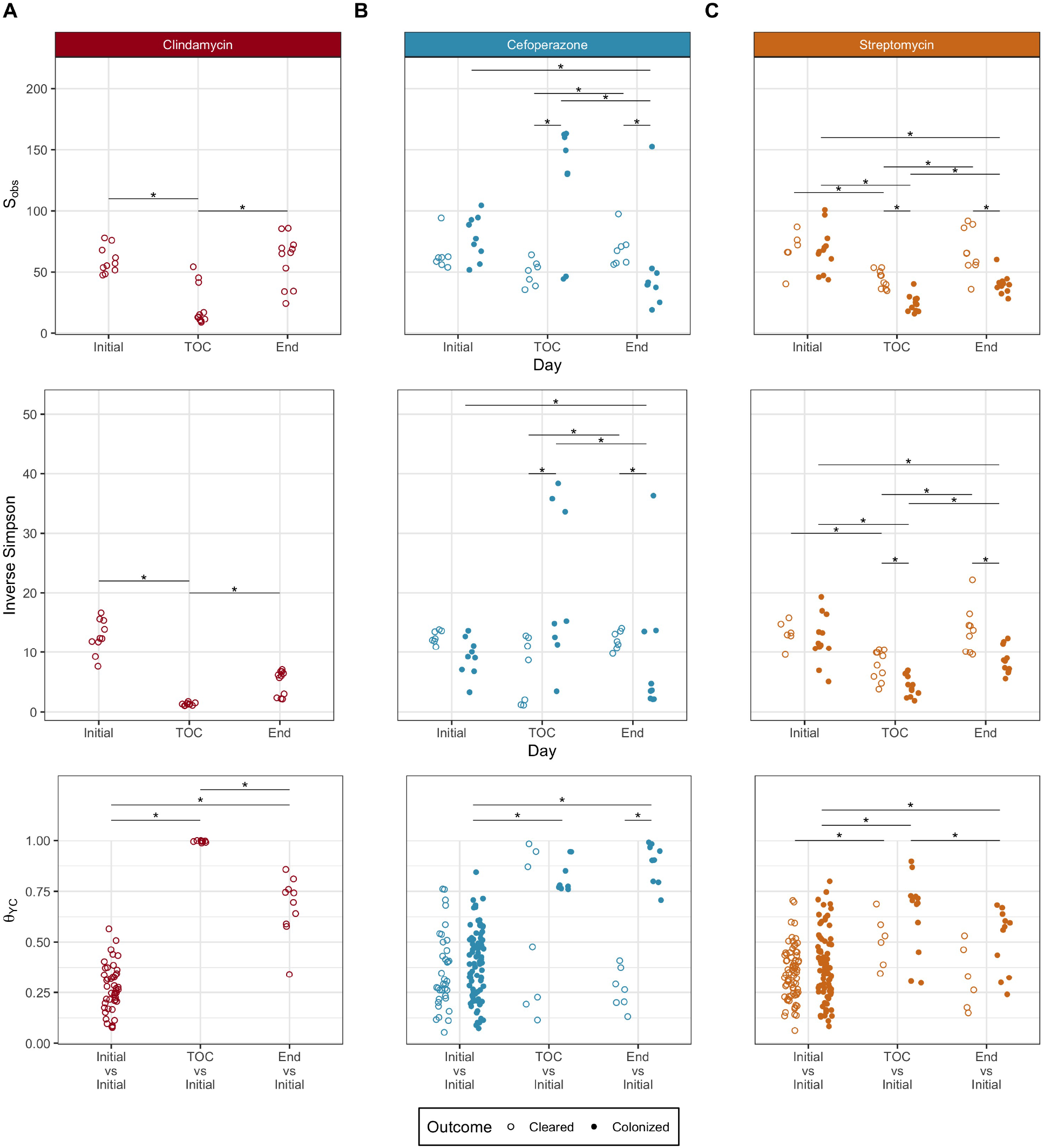
Microbiota community diversity showed antibiotic-specific trends associated with *C. difficle* colonization clearance. For communities colonized with *C. difficile* from mice treated with clindamycin (A), cefoperazone (B), and streptomycin (C), microbiota *α*-diversity (S_obs_ and Inverse Simpson) and *β*-diversity (*θ*_YC_) were compared at the initial pre-antibiotic treatment state, time of *C. difficile* challenge (TOC), and end of the experiment. *β*-diversity (*θ*_YC_) was compared between the initial pre-antibiotic treatment to all other initial pre-antibiotic treatment communities treated with the same antibiotic, the initial community to the same community at the time of *C. difficile* challenge, and the initial community to the same community at end of the experiment. (clindamycin - cleared N = 11; cefoperazone - cleared N = 7, colonized N = 9; streptomycin - cleared N = 9, colonized N = 11). * indicates statistical significance of *P* < 0.05, calculated by Wilcoxon rank sum test with Benjamini-Hochberg correction.

We applied the same analysis to the cefoperazone-treated mice to understand what community features were relevant to clearing *C. difficile.* Increasing the dose of cefoperazone shifted the dominant community members from relatives of the *Porphyromonadaceae, Bacteroides* and *Akkermansia* to relatives of the *Lactobacillus* and *Enterobacteriaceae* at the time of challenge (Figure 1E, S1). We saw a similar increase in relatives of *Enterobacteriaceae* with clindamycin. However, the cefoperazone-treated mice that had larger increases in *Enterobacteriaceae* were unable to clear *C. difficile.* We next investigated the differences between the cefoperazone-treated mice that cleared *C. difficile* to those that did not. For the communities that cleared *C.difficile*,diversity was maintained throughout the experiment (Figure 2B). The mice treated with cefoperazone that remained colonized experienced an increase in alpha diversity, likely driven by the decrease in highly abundant populations and increase in low abundant populations (Figure 1E). These persistently colonized communities also had a large shift away from the initial community structure caused by the antibiotic treatment (P < 0.05), which remained through the end of the experiment (P < 0.05) (Figure 2B). These data suggested that it was necessary for cefoperazone-treated mice to become more similar to the initial pre-antibiotic community structure to clear *C. difficile.* We next investigated the changes in OTU abundances between the communities that cleared *C. difficile* and those that did not to elucidate the community members involved in clearance. Communities that remained colonized were significantly enriched in facultative anaerobic populations including *Enterococcus, Pseudomonas, Staphylococcus*, and *Enterobacteriaceae* at the time of challenge. Communities that cleared *C. difficile* had significant enrichment in 10 different OTUs related to the *Porphyromonadaceae* at the end of the experiment (Figure 3A). We were also interested in the temporal changes within each community so we investigated which OTUs changed due to antibiotic treatment or during the *C. difficile* colonization. The majority of significant temporal differences in OTUs for cefoperazone-treated mice occurred in persistently colonized communities. Persistently colonized communities had a persistent loss of numerous relatives of the *Porphyromonadaceae* and increases in the relative abundance of facultative anaerobes (Figure 4C, S2). Overall, persistent *C. difficile* colonization in cefoperazone-treated mice was associated with a shift in the microbiota to a new community structure which seemed unable to recover from the antibiotic perturbation, whereas clearance occurred when the community was capable of returning to its original structure.

**Figure 3.**
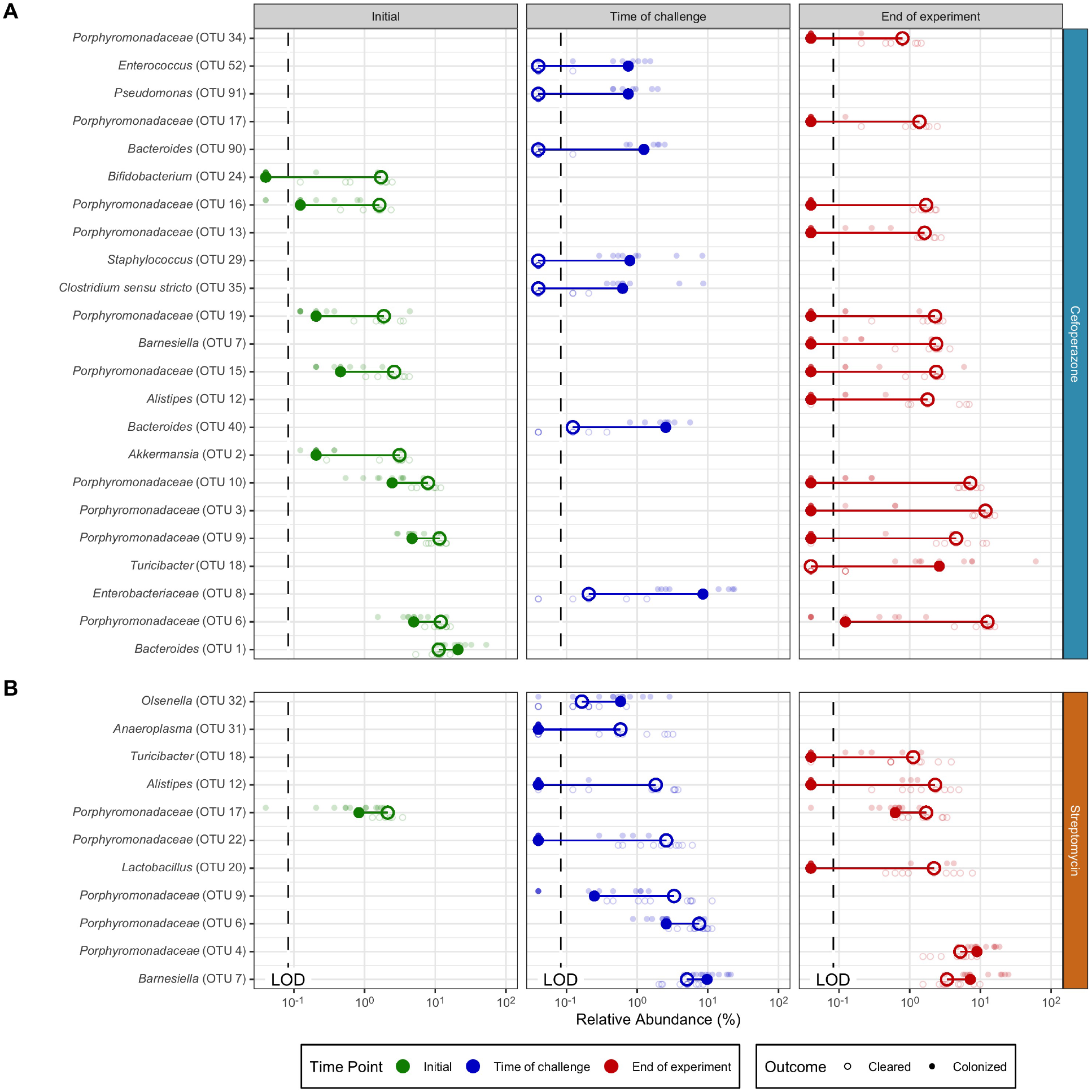
OTU abundance differences between communities that cleared *C. difficile* colonization and remained colonized are unique to each treatment. For cefoperazone (A) and streptomycin (B), the difference in the relative abundance of OTUs that were significantly different between communities that eliminated *C. difficile* colonization and those that remained colonized within each antibiotic treatment for each time point. Bold points are median relative abundance and transparent points are relative abundance of individual mice. Lines connect points within each comparison to show difference in medians. Only OTUs at time points with statistically significant differences, *P* < 0.05, were plotted (calculated by Wilcoxon rank sum test with Benjamini-Hochberg correction). Limit of detection (LOD).

**Figure 4.**
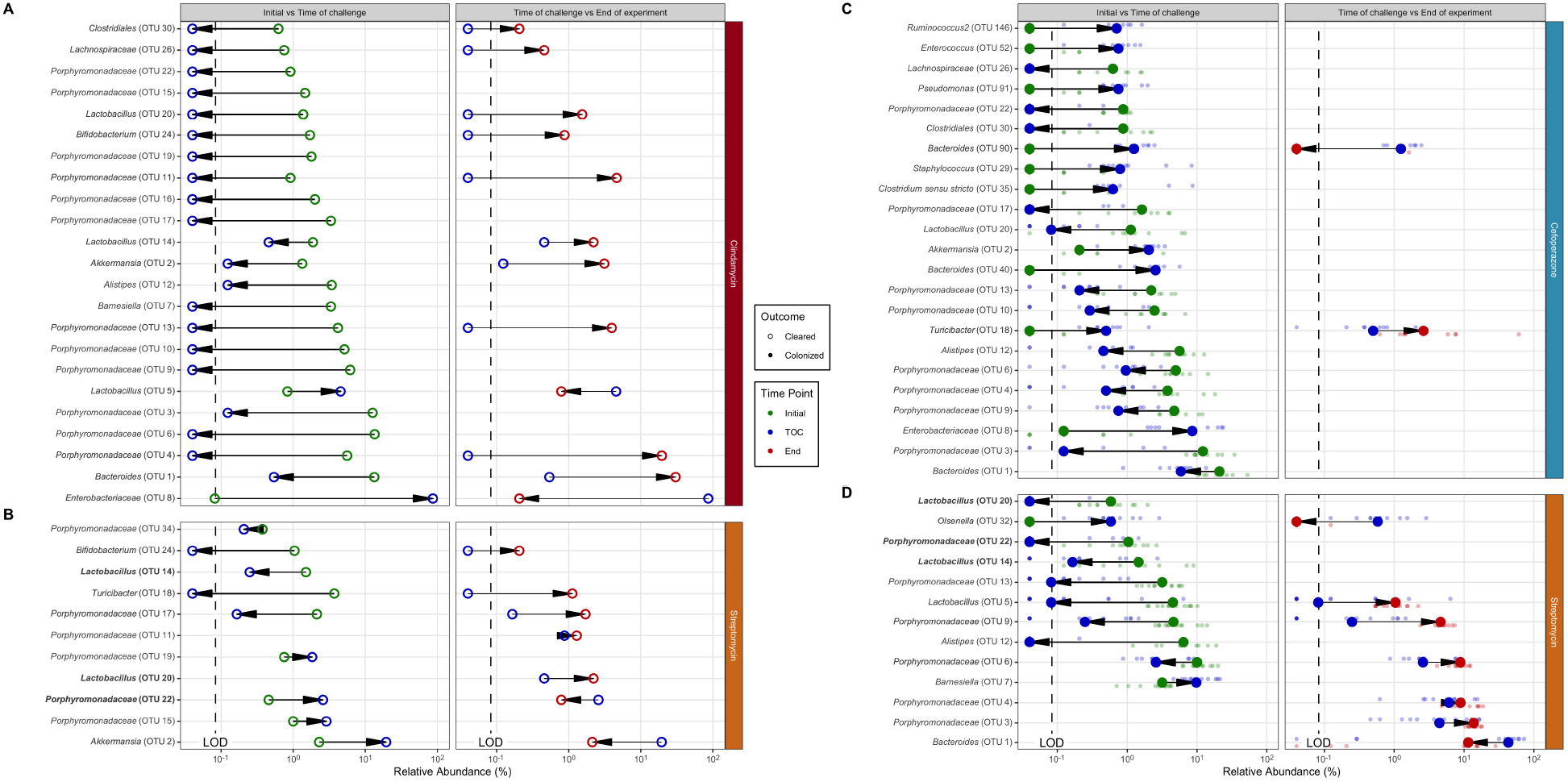
Each antibiotic had specific sets of temporal changes in OTU abundance associated with *C. difficile* colonization and clearance. For clindamycin (A), cefoperazone (C), and streptomycin (B, D), the difference in the relative abundance of OTUs that were significantly different between time points within each *C. difficile* colonization outcome for each antibiotic treatment. Bold points are median relative abundance and transparent points are relative abundance of individual mice. Lines connect points within each comparison to show difference in medians. Arrows point in the direction of the temporal change of the relative abundance. Only OTUs at time points with statistically significant differences, *P* < 0.05, were plotted (calculated by Wilcoxon rank sum test with Benjamini-Hochberg correction). Bold OTUs were shared across outcomes. Limit of detection (LOD).

Finally, we identified the differences in *C. difficile* colonization for streptomycin-treated mice. Increasing the dose of streptomycin maintained the abundance of relatives of the *Porphyromonadaceae* and *Bacteroides*, but reduced most of the other genera including populations of the *Lactobacillus, Lachnospiraceae, Ruminococcaceae, Alistipes*, and *Clostridiales* (Figure 1F). Both communities that cleared and those that remained colonized had similar changes in diversity. Streptomycin-treated mice became mildly dissimilar (P < 0.05) and less diverse (P < 0.05) with streptomycin treatment but by the end of the experiment returned to resemble the pre-antibiotic community (P < 0.05) (Figure 2C). Those communities that remained colonized had slightly lower alpha-diversity than those that cleared *C. difficile. (P* < 0.05). Persistently colonized mice had reduced relative abundance of relatives of *Alistipes, Anaeroplasma*, and *Porphyromonadaceae* at time of challenge compared to the mice that cleared *C. difficle* (Figure 3B). At the end of the experiment the mice that were still colonized had lower abundances of *Turicibacter, Alistipes*, and *Lactobacillus.* Since most of the differences were reduced relative abundances in the colonized mice, we were interested to explore what temporal changes occurred between pre-antibiotic treatment, the time of challenge, and the end of the experiment for the communities that cleared *C. difficile.* The temporal changes in streptomycin-treated mice were more subtle than those observed with the other antibiotic treatments. At the time of challenge, the communities that remained colonization had reductions in 4 OTUs related to the *Porphyromonadaceae.* Those that cleared *C. difficile* also had changes in OTUs related to the *Porphyromonadaceae*, however, 2 populations decreased and 2 increased in abundance (Figure 4B, D). At the end of the experiment, all communities experienced recovery of the abundance of many of the populations changed by the streptomycin treatment, but the communities that remained colonized did not recover 5 of the OTUs of *Alistipes*, *Lactobacillus*, and *Porphyromonadaceae* that were reduced by streptomycin. The differences between the streptomycin-treated mice that remained colonized and those had been cleared of *C. difficile* were not as distinct as those observed with the cefoperazone treatment. The differences between colonized and cleared streptomycin-treated mice were minimal, which suggested the few differences may be responsible for the clearance. Overall, these data revealed that while there were commonly affected families across the antibiotic treatments, such as the *Porphyromonadaceae, C. difficile* clearance was associated with community and OTU differences specific to each antibiotic.

### Distinct features of the bacterial community at the time of infection predicted end point colonization

To determine whether the community composition at the time of *C. difficile* challenge could predict *C. difficile* clearance, we built a machine learning model using L2 logistic regression. We evaluated the predictive performance of the model using the area under the receiver operating characteristic curve (AUROC), where a value of 0.5 indicated the model is random and 1.0 indicated the model always correctly predicts the outcome. Our model resulted in a AUROC of 0.986 [IQR 0.970-1.000], which suggested that the model was able to use the relative abundance of OTUs at the time of challenge to accurately predict colonization clearance (Figure S3). To assess the important features, we randomly permuted each OTU feature by removing it from the training set to determine its effect on the prediction (Figure 5A). The most important feature was an OTU related to the *Enterobacteriaceae*, whose abundance predicted clearance. This result appears to have been strongly driven by the clindamycin data (Figure 5B, C). The remaining OTU features did not have a large effect on the model performance, which suggested that the model decision was spread across many features. These results revealed the model used the relative abundance data of the community members and the relationship between those abundances to correctly classify clearance. There were many OTUs with treatment and outcome specific abundance patterns that did not agree with the odds ratio of the OTU used by the model. For example, *Enterobacteriaceae* abundance influenced the model to predict clearance (Figure 5B), however in experiments that used cefoperazone, the communities that remained colonized had higher abundances of *Enterobacteriaceae* than the communities that cleared colonization (Figure 5C). The model arrived at the correct prediction through the influence of other OTUs. Therefore, the model used different combinations of multiple OTUs and their relative abundances across treatments to predict *C. difficile* clearance. These data can offer a basis for hypotheses regarding the distinct combinations of bacteria that promote *C. difficile* clearance.

**Figure 5.**
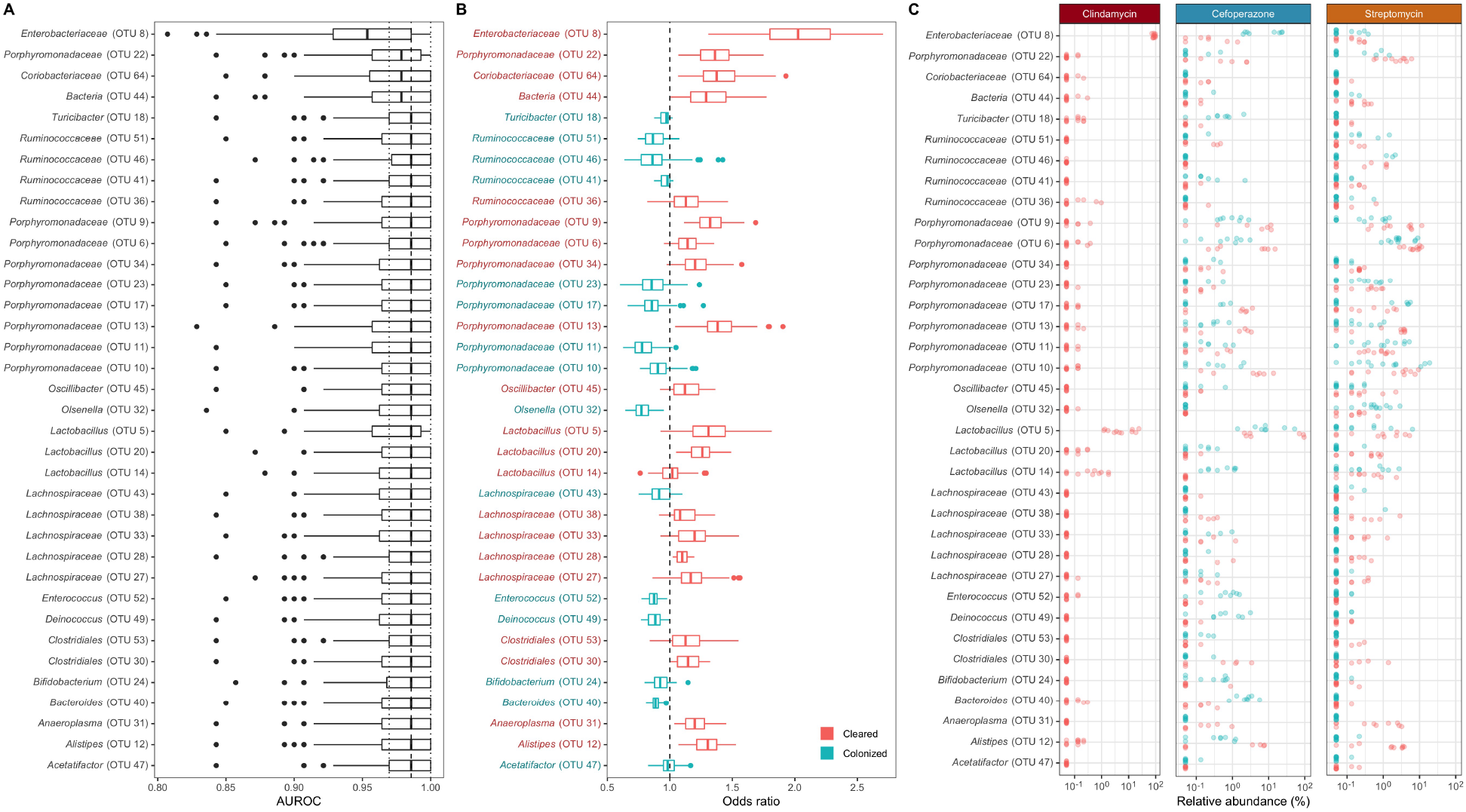
Distinct features of the bacterial community at the time of infection can classify end point colonization. (A) L2 logistic regression model features’ importance determined by the decrease in model performance when randomizing an individual feature. All OTUs affecting performance shown. Dashed lines show performance range of final model with all features included. (B) Distribution of odds ratio used in L2 logistic regression model. Values above 1 indicate abundance predicted the community cleared colonization (red) and values below 1 indicate abundance predicted *C. difficile* remained colonized (blue). Feature label and boxplot are colored to match the median odds ratio. (C) Relative abundance difference in features used by L2 logistic regression model displayed by antibiotic treatment.

### Conditional independence networks revealed treatment-specific relationships between the community members and C. *difficile* during colonization clearance

We next investigated the relationship between temporal changes in the community and *C. difficile* by building a conditional independence network for each treatment using SPIEC-EASI (sparse inverse covariance estimation for ecological association inference) (21). First, we focused on the first-order associations of *C. difficile* (Figure 6A). In clindamycin-treated mice, *C. difficile* had positive associations with relatives of *Enterobacteriaceae*, *Pseudomonas*, and *Olsenella* and negative associations with relatives of the *Lachnospiraceae* and *Clostridium* XIVa. *C. difficile* had limited associations in cefoperazone-treated mice; the primary association was positive with relatives of *Enterobacteriaceae.* In streptomycin-treated mice, *C. difficile* had negative associations with relatives of the *Porphyromonadaceae* and positive associations with populations of the *Ruminococcaceae, Bacteroidetes, Clostridium* IV and *Olsenella.* Next, we quantified the degree centrality, the number of associations between each OTU for the whole network of each antibiotic and outcome, and betweenness centrality, the number of associations connecting two OTUs that pass through an OTU (Figure 6B). This analysis revealed cefoperazone treatment resulted in networks primarily composed of singular associations with much lower degree centrality (P < 0.05) and betweenness centrality (P <0.05) than the other antibiotic treatments. Communities that were treated with cefoperazone that resulted in cleared or persistent colonization had 10 to 100-fold lower betweenness centrality values than communities treated with clindamycin or streptomycin. Collectively, these networks suggest *C. difficile* colonization was affected by unique sets of OTUs in mice treated with clindamycin and streptomycin, but cefoperazone treatment eliminated bacteria critical to maintaining community interactions and had few populations that associated with *C. difficile*.

**Figure 6.**
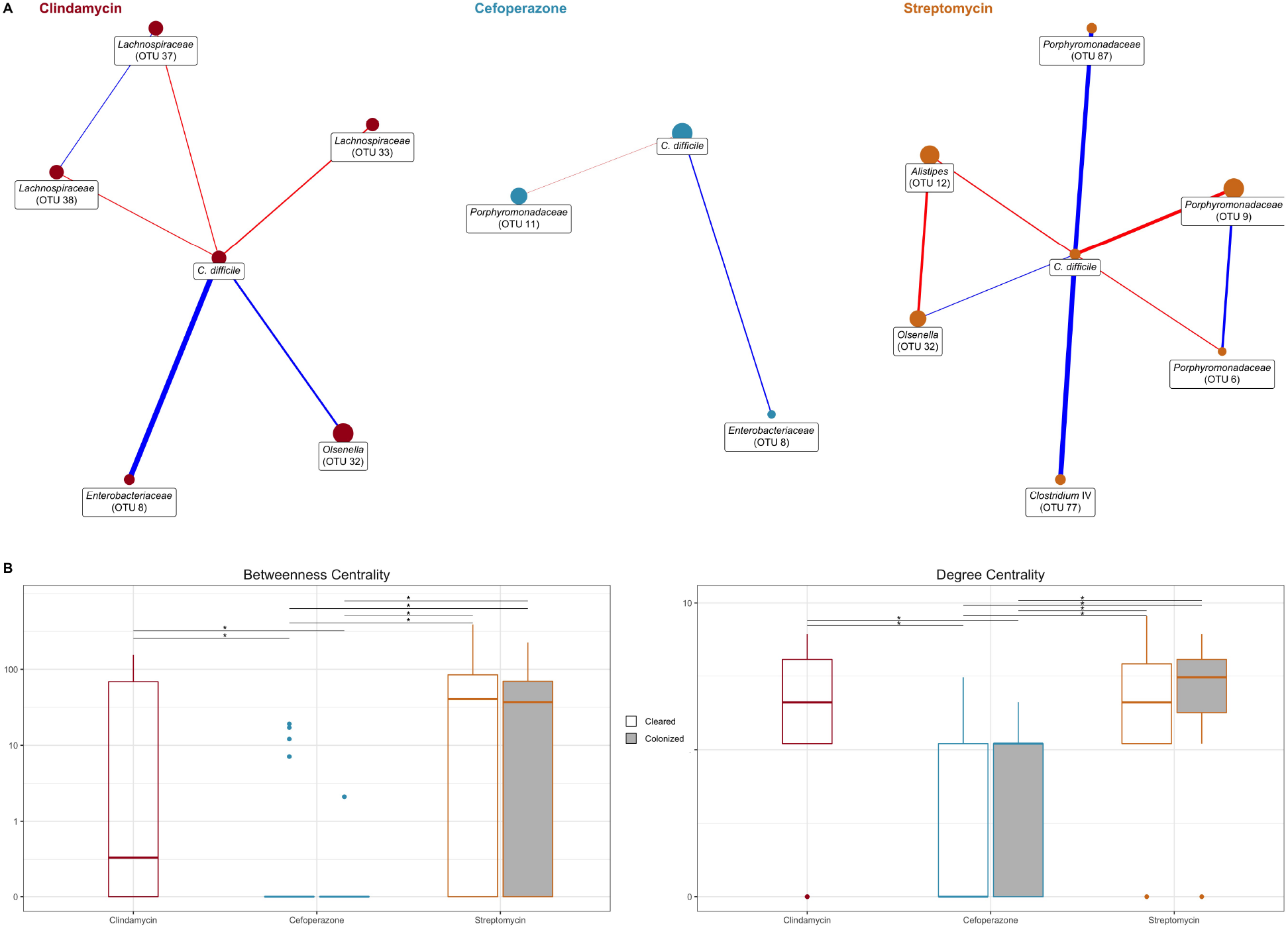
Conditional independence networks reveal treatment-specific relationships between the community and *C. difficile* during colonization clearance. (A) SPIEC-EASI (sparse inverse covariance estimation for ecological association inference) networks showing conditionally independent first-order relationships between *C. difficile* and the community as *C. difficile* was cleared from the gut environment. Nodes are sized by median relative abundance of the OTU. A red colored edge indicates a negative interaction and blue indicates a positive interaction, while edge thickness indicates the interaction strength. (B) Network centrality measured with betweenness, i.e. how many paths between two OTUs pass through an individual, and degree, i.e. how many connections an OTU had. * indicates statistical significance of *P* < 0.05, calculated by Wilcoxon rank sum test with Benjamini-Hochberg correction.

## Discussion

We have shown that different antibiotic treatments resulted in specific changes to the microbiota that were associated with *C. difficile* clearance. Clindamycin-treated mice became susceptible with a dominant bloom in populations related to *Enterobacteriaceae.* Clearance was associated with the resolution of the bloom and recovery of bacteria that were reduced by the antibiotic treatment. Cefoperazone-treated mice became susceptible with the expansion of numerous facultative anaerobes. Communities with a sustained presence of these facultative anaerobes were unable to recover from the initial antibiotic perturbation or clear the colonization, whereas the communities that returned to their initial community were able to clear *C. difficile* colonization. Streptomycin-treated mice became susceptible with fewer and smaller changes than the other treatments. The communities that cleared colonization had slightly higher *α*-diversity than those that remained colonized. Additionally, all communities in mice treated with streptomycin had similar numbers of OTUs changing through the experiment but the specific OTUs were different for each outcome. These observations support our hypothesis that each colonized community has antibiotic-specific changes that create unique conditions for *C. difficile* colonization and requires specific changes within each community to clear *C. difficile.*

Previous studies have identified microbiota associated with *C. difficile* colonization resistance in either a set of closely related murine communities or collectively across many different susceptible communities (11, 15, 22). These bacteria were then tested in *C. difficile* infection models. These experiments were able to show decreased colonization but were unable to fully clear *C. difficile*(11, 23). Rather than looking for similarities across all susceptible communities, we explored the changes that were associated with *C. difficile* clearance for each antibiotic. Even though these mice all came from the same breeding colony with similar initial microbiomes, *C. difficile* clearance was associated with antibiotic-specific changes in community diversity, OTU abundances, and associations between OTUs. Our data suggest that the set of bacteria necessary to restore colonization resistance following one antibiotic perturbation may not be effective for all antibiotic perturbations. We have developed this modeling framework starting from a single mouse community. It should also be relevant when considering interpersonal variation among humans (24).

Recent studies have begun to uncover how communities affect *C. difficile* colonization (17–20, 24). We attempted to understand the general trends in each antibiotic treatment that lead to clearance of *C. difficile*. We categorized the general changes and microbial relationships of these experiments into three models. First, a model of temporary opportunity characterized by the transient dominance of a facultative anaerobe which permits *C. difficile* colonization but *C. difficile* is not able to persist, as with clindamycin treatment. We hypothesize this susceptibility is due to a transient repression of community members and interventions which further perturb the community may worsen the infection. Time alone may be sufficient for the community to clear colonization (15, 22, 25) but treating the community with an antibiotic or the bowel preparation for an FMT (26, 27) may prolong susceptibility by eliminating protective functions or opening new niches. Second, a model of an extensive opportunity characterized by a significant perturbation leading to a persistent increase in facultative anaerobes and exposing multiple niches, as with cefoperazone treatment. These communities appear to have been severely depleted of multiple critical community members and are likely lacking numerous protective functions (20). We hypothesize multiple niches are available for *C. difficile* to colonize. In this scenario, a full FMT may be insufficient to provide adequate diversity and abundance to outcompete and occupy all the exposed niches. Multiple FMTs (28, 29) or transplant of an enriched fecal community (30) may be necessary to recover the microbiota enough to outcompete *C. difficile* for the nutrient niches and replace the missing protective functions. Third, a model of a specific opportunity characterized by a perturbation that only affects a select portion of the microbiota, leading to small changes in relative abundance and a slight decrease in diversity, opening a limited niche for *C. difficile* to colonize, as with streptomycin treatment. We hypothesize that a few specific bacteria would be necessary to recolonize the exposed niche space and eliminate *C. difficile* colonization (13, 17). A fecal microbiota transplant may contain the bacterial diversity needed to fill the open niche space and help supplant *C. difficile* from the exposed niche of the colonized community. Analyzing each of these colonization models individually allowed us to understand how each may clear *C. difficile* colonization.

Future investigations can further identify the exposed niches of susceptible communities and the requirements to clear *C. difficile* colonization. One common theme for susceptibility across treatments was the increased abundance of facultative anaerobes. These blooms of facultative anaerobes could be attributed to the loss of the indigenous obligate anaerobes with antibiotic treatment (31, 32). However, it is unclear what prevents the succession from the facultative anaerobes back to the obligate anaerobes in cefoperazone-treated mice. Future studies should investigate the relationship between facultative anaerobe blooms and susceptibility to colonization as well as interventions to recover the obligate anaerobes. Another aspect to consider in future experiments is *C. difficile* strain specificity. Other strains may fill different niche space and fill other community interactions (33–35). For example, more virulent strains, like *C. difficile* VPI 10463, may have a greater effect on the gut environment since it produces more toxin (15, 36). Those differences could have different impacts on the susceptible community and change the requirements to clear *C. difficile.* Finally, we have shown that the functions found in communities at peak colonization are antibiotic-specific (20). Here, we have shown the community changes associated with *C. difficile* clearance are antibiotic-specific. It is unknown how the community functions contributing to *C. difficile* clearance compare across antibiotics. Examining the changes in transcription and metabolites during clearance will help define the activities necessary to clear *C. difficile* and if they are specific to the perturbation. This information will build upon the community differences presented in this study and move us closer to elucidating how the microbiota clears *C. difficile* colonization and developing targeted therapeutics.

We have shown that mice became susceptible to *C. difficile* colonization after three different antibiotic treatments and then differed in their ability to clear the colonization. These experiments have shown that each antibiotic treatment resulted in different community changes leading to *C. difficile* clearance. These differences suggest that a single mechanism of infection and one treatment for all *C. difficile* infections may not be appropriate. While our current use of FMT to eliminate CDI is highly effective, it does not work in all patients and has even resulted in adverse consequences (7–10). The findings in this study may help explain why FMTs may be ineffective. Although an FMT transplants a whole community, it may not be sufficient to replace the missing community members or functions to clear *C. difficile.* Alternatively, the FMT procedure itself may disrupt the natural recovery of the community. The knowledge of how a community clears *C. difficile* colonization will advance our ability to develop targeted therapies to manage CDI.

## Materials and Methods

### Animal care

All mice were obtained from a single breeding colony and maintained in specific-pathogen-free (SPF) conditions at the University of Michigan animal facility. All mouse protocols and experiments were approved by the University Committee on Use and Care of Animals at the University of Michigan and completed in agreement with approved guidelines.

### Antibiotic administration

Mice were given one of three antibiotics, cefoperazone, clindamycin, or streptomycin. Cefoperazone (0.5, 0.3, or 0.1 mg/ml) and streptomycin (5, 0.5, or 0.1 mg/ml) were delivered via drinking water for 5 days. Clindamycin (10 mg/kg) was administered through intraperitoneal injection.

### C. *difficile* challenge

Mice were returned to untreated drinking water for 24 hours before challenging with *C. difficile* strain 630Δerm spores. *C. difficile* spores were aliquoted from a single spore stock stored at 4°C. Spore concentration was determined one week prior to the day of challenge (37). 10^3^ *C. difficile* spores were orally gavaged into each mouse. Once the gavages were completed, the remaining spore solution was serially diluted and plated to confirm the spore concentration that was delivered.

### Sample collection

Fecal samples were collected on the day antibiotic treatment was started, on the day of *C. difficile* challenge and the following 10 days. For the day of challenge and beyond, a fecal sample was also collected and weighed. Under anaerobic conditions a fecal sample was serially diluted in anaerobic phosphate-buffered saline and plated on TCCFA plates. After 24 hours of anaerobic incubation at 37°C, the number of colony forming units (CFU) were determined (38).

### DNA sequencing

Total bacterial DNA was extracted from each fecal sample using MOBIO PowerSoil-htp 96-well soil DNA isolation kit. We created amplicons of the 16S rRNA gene V4 region and sequenced them using an Illumina MiSeq as described previously (39).

### Sequence curation

Sequences were processed using mothur(v.1.43.0) as previously described (39). Briefly, we used a 3% dissimilarity cutoff to group sequences into operational taxonomic units (OTUs). We used a naive Bayesian classifier with the Ribosomal Database Project training set (version 16) to assign taxonomic classifications to each OTU (41). With the fecal samples, we also sequenced a mock community with a known community composition and their true 16s rRNA gene sequences. We processed this mock community along with our samples to determine our sequence curation resulted in an error rate of 0.019%.

### Statistical analysis and modeling

Diversity comparisons were calculated in mothur. To compare *α*-diversity metrics, we calculated the number of OTUs (S_obs_) and the Inverse Simpson diversity index. To compare across communities, we calculated dissimilarity matrices based on metric of Yue and Clayton (42). All calculations were made by rarifying samples to 1,200 sequences per sample to limit biases due to uneven sampling. OTUs were subsampled to 1,200 counts per sample and remaining statistical analysis and data visualization was performed in R (v3.5.1) with the tidyverse package (v1.3.0). Significance of pairwise comparisons of *α*-diversity (S_obs_ and Inverse Simpson), *β*-diversity (*θ*_YC_), OTU abundance, and network centrality (betweenness and degree) were calculated by pairwise Wilcoxon rank sum test and then *P* values were corrected for multiple comparisons with a Benjamini and Hochberg adjustment for a type I error rate of 0.05 (43). Logistic regression models were constructed with OTUs from all day 0 samples using half of the samples to train and the other half to test the model. The model was developed from the caret R package (v6.0-85) and previously developed machine learning pipeline (44). For each antibiotic treatment, conditional independence networks were calculated from the day 1 through 10 samples of all mice initially colonized using SPIEC-EASI (sparse inverse covariance estimation for ecological association inference) methods from the SpiecEasi R package after optimizing lambda to 0.001 with a network stability between 0.045 and 0.05 (v1.0.7) (21). Network centrality measures degree and betweenness were calculated on whole networks using functions from the igraph R package (v1.2.4.1).

### Code availability

Scripts necessary to reproduce our analysis and this paper are available in an online repository (https://github.com/SchlossLab/Lesniak_Clearance_XXXX_2020).

### Sequence data accession number

All 16S rRNA gene sequence data and associated metadata are available through the Sequence Read Archive via accession PRJNA674858.

## Acknowledgements

Thank you to Begüm Topçuoglu and Sarah Tomkovich for critical discussion in the development and execution of this project. This work was supported by several grants from the National Institutes for Health R01GM099514, U19AI090871, U01AI12455, and P30DK034933. Additionally, NAL was supported by the Molecular Mechanisms of Microbial Pathogenesis training grant (NIH T32 AI007528). The funding agencies had no role in study design, data collection and analysis, decision to publish, or preparation of the manuscript.

---

**Figure S1.**
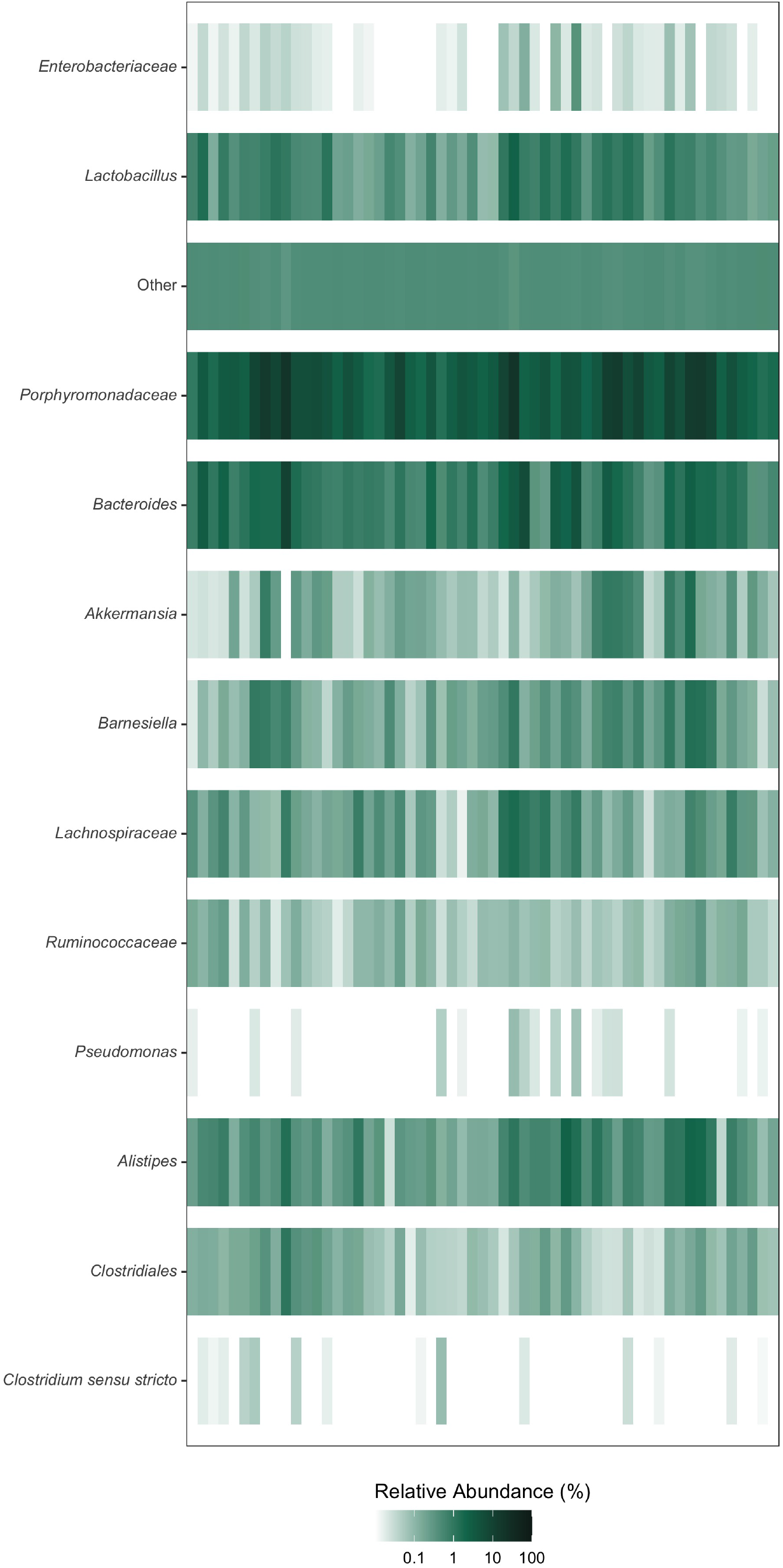
Initial microbiota relative abundance of mice prior to antibiotic treatment. Relative abundance at the beginning of the experiment prior to antibiotic treatment of twelve most abundant genera post antibiotic treatment, all other genera grouped into Other. Each column is an individual mouse. Color intensity is log_10_-transformed mean percent relative abundance of each day. (N = 57).

**Figure S2.**
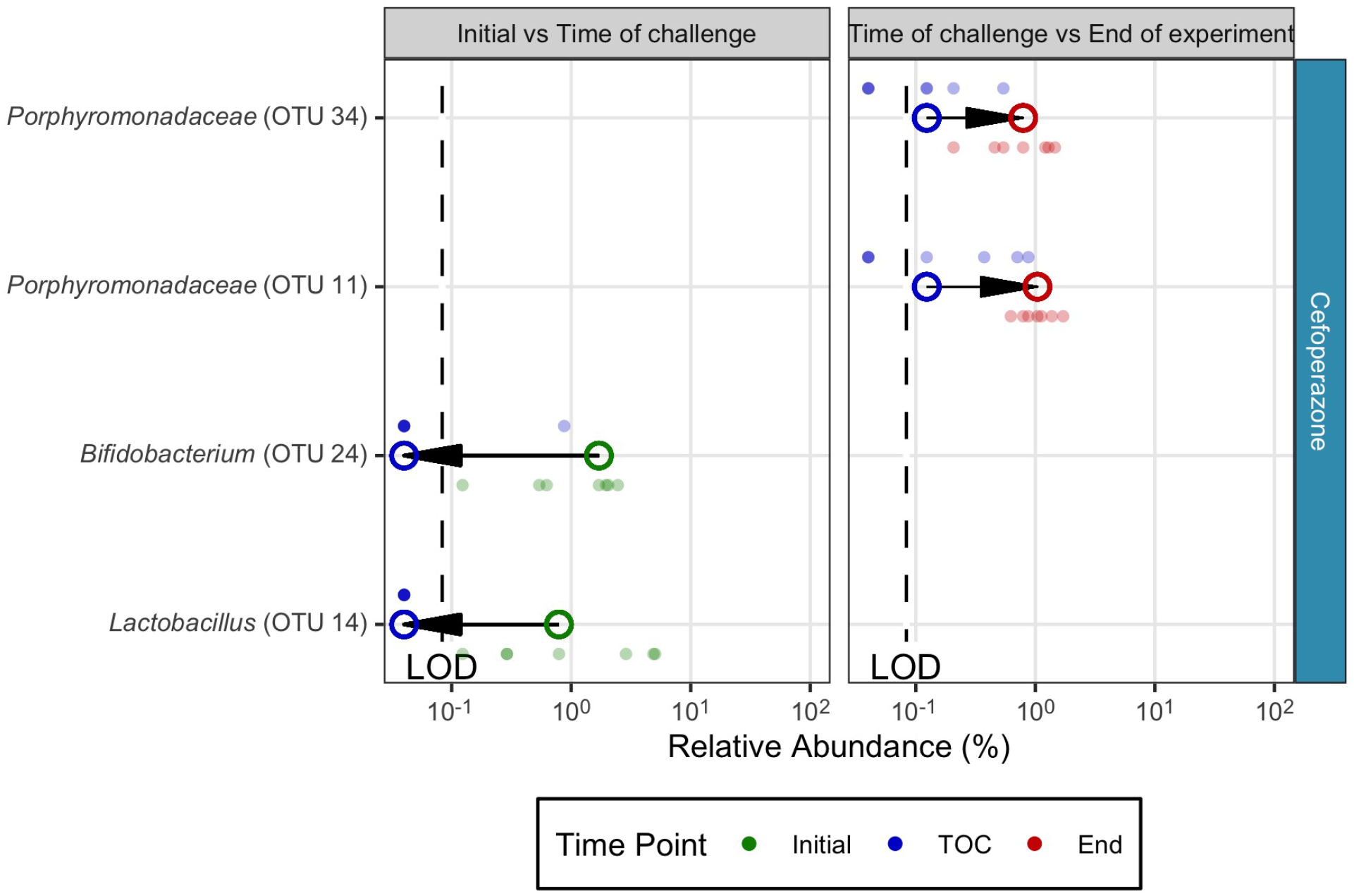
Temporally differing OTU for cefoperazone-treated mice that cleared *C. difficile* colonization. Bold points are median relative abundance and transparent points are relative abundance of individual mice. Lines connect points within each comparison to show difference in medians. Arrows point in the direction of the temporal change of the relative abundance. Only OTUs at time points with statistically significant differences, *P* < 0.05, were plotted (calculated by Wilcoxon rank sum test with Benjamini-Hochberg correction). Limit of detection (LOD).

**Figure S3.**
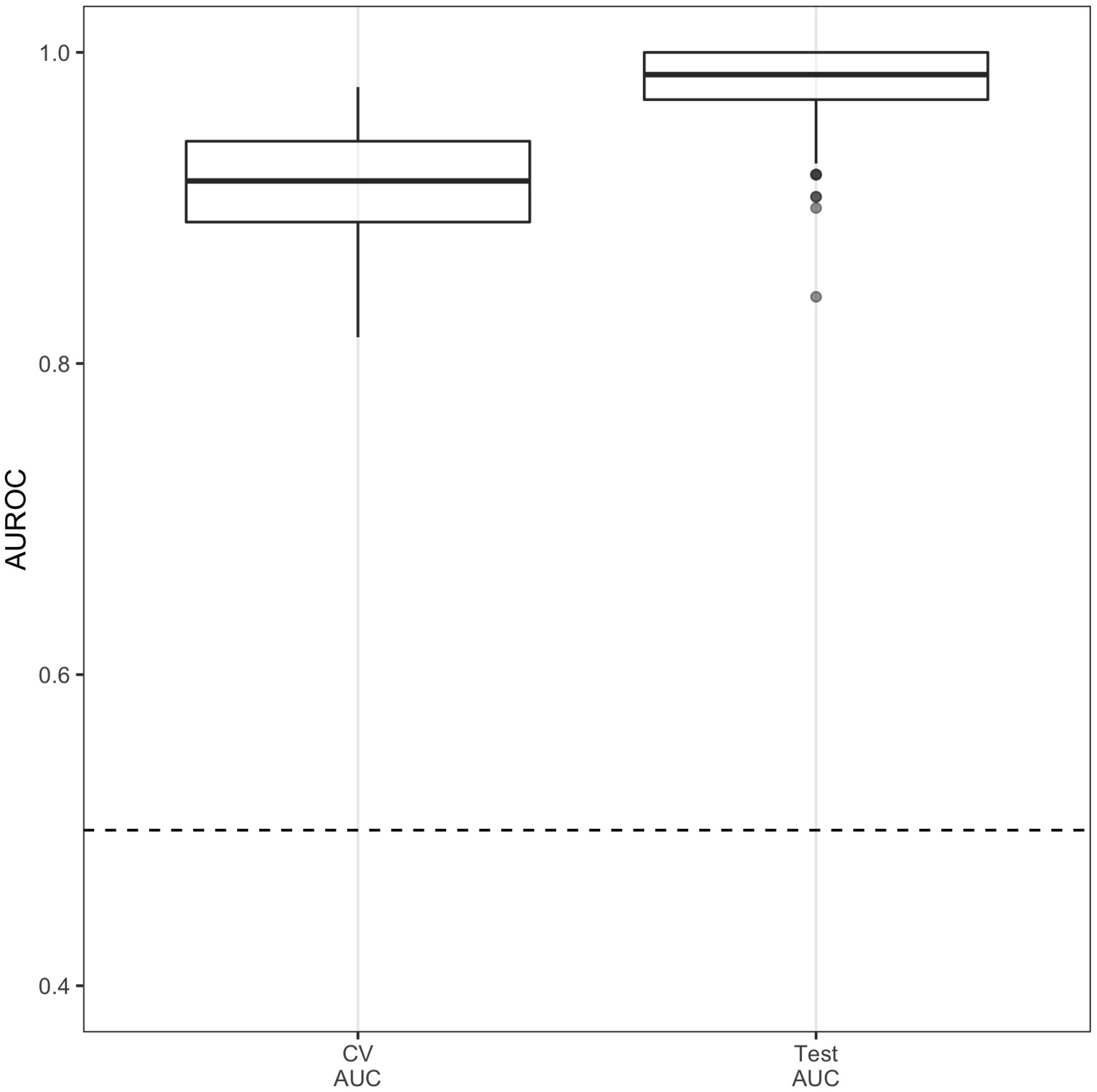
Bacterial community at the time of infection can classify endpoint colonization. Classification performance of L2 logistic regression. Area under the receiver-operator curve for classifying if the community will remain colonized based on the OTUs present at the time of *C. difficile* infection (Day 0). Cross-validation of model performed on half of the data to tune model (CV AUC) and then tuned model was tested on the held-out data (Test AUC).

## References

1. Ducarmon QR, Zwittink RD, Hornung BVH, Schaik W van, Young VB, Kuijper EJ. 2019. Gut microbiota and colonization resistance against bacterial enteric infection. Microbiology and Molecular Biology Reviews 83. doi:10.1128/mmbr.00007-19.

2. Britton RA, Young VB. 2012. Interaction between the intestinal microbiota and host in clostridium difficile colonization resistance. Trends in Microbiology 20:313–319. doi:10.1016/j.tim.2012.04.001.

3. Lessa FC, Mu Y, Bamberg WM, Beldavs ZG, Dumyati GK, Dunn JR, Farley MM, Holzbauer SM, Meek JI, Phipps EC, Wilson LE, Winston LG, Cohen JA, Limbago BM, Fridkin SK, Gerding DN, McDonald LC. 2015. Burden of Clostridium difficile Infection in the united states. New England Journal of Medicine 372:825–834. doi:10.1056/nejmoa1408913.

4. Zimlichman E, Henderson D, Tamir O, Franz C, Song P, Yamin CK, Keohane C, Denham CR, Bates DW. 2013. Health careAssociated infections. JAMA Internal Medicine 173:2039. doi:10.1001/jamainternmed.2013.9763.

5. Spigaglia P, Barbanti F, Morandi M, Moro ML, Mastrantonio P. 2016. Diagnostic testing for clostridium difficile in italian microbiological laboratories. Anaerobe 37:29–33. doi:10.1016/j.anaerobe.2015.11.002.

6. Dieterle MG, Rao K, Young VB. 2018. Novel therapies and preventative strategies for primary and recurrent Clostridium difficile infections. Annals of the New York Academy of Sciences 1435:110–138. doi:10.1111/nyas.13958.

7. Juul FE, Garborg K, Bretthauer M, Skudal H, Øines MN, Wiig H, Rose, Seip B, Lamont JT, Midtvedt T, Valeur J, Kalager M, Holme, Helsingen L, Løberg M, Adami H-O. 2018. Fecal microbiota transplantation for primary clostridium difficile infection. New England Journal of Medicine 378:2535–2536. doi:10.1056/nejmc1803103.

8. Seekatz AM, Aas J, Gessert CE, Rubin TA, Saman DM, Bakken JS, Young VB. 2014. Recovery of the gut microbiome following fecal microbiota transplantation. mBio 5. doi:10.1128/mbio.00893-14.

9. Patron RL, Hartmann CA, Allen S, Griesbach CL, Kosiorek HE, DiBaise JK, Orenstein R. 2017. Vancomycin taper and risk of failure of fecal microbiota transplantation in patients with recurrent clostridium difficile infection. Clinical Infectious Diseases 65:1214–1217. doi:10.1093/cid/cix511.

10. DeFilipp Z, Bloom PP, Soto MT, Mansour MK, Sater MRA, Huntley MH, TurbettS, Chung RT, Chen Y-B, Hohmann EL. 2019. Drug-resistant e. Coli bacteremia transmitted by fecal microbiota transplant. New England Journal of Medicine 381:2043–2050. doi:10.1056/nejmoa1910437.

11. Buffie CG, Bucci V, Stein RR, McKenney PT, Ling L, Gobourne A, No D, Liu H, Kinnebrew M, Viale A, Littmann E, Brink MRM van den, Jenq RR, Taur Y, Sander C, Cross JR, Toussaint NC, Xavier JB, Pamer EG. 2014. Precision microbiome reconstitution restores bile acid mediated resistance to clostridium difficile. Nature 517:205–208. doi:10.1038/nature13828.

12. Fletcher JR, Erwin S, Lanzas C, Theriot CM. 2018. Shifts in the gut metabolome and clostridium difficile transcriptome throughout colonization and infection in a mouse model. mSphere 3. doi:10.1128/msphere.00089-18.

13. Reed AD, Nethery MA, Stewart A, Barrangou R, Theriot CM. 2020. Strain-dependent inhibition of clostridioides difficile by commensal clostridia carrying the bile acid-inducible (bai) operon. Journal of Bacteriology 202. doi:10.1128/jb.00039-20.

14. Jenior ML, Leslie JL, Young VB, Schloss PD. 2017. Clostridium difficile colonizes alternative nutrient niches during infection across distinct murine gut microbiomes. mSystems 2. doi:10.1128/msystems.00063-17.

15. Lawley TD, Clare S, Walker AW, Stares MD, Connor TR, Raisen C, Goulding D, Rad R, Schreiber F, Brandt C, Deakin LJ, Pickard DJ, Duncan SH, Flint HJ, Clark TG, Parkhill J, Dougan G. 2012. Targeted restoration of the intestinal microbiota with a simple, defined bacteriotherapy resolves relapsing clostridium difficile disease in mice. PLoS Pathogens 8:e1002995. doi:10.1371/journal.ppat.1002995.

16. McDonald JAK, Mullish BH, Pechlivanis A, Liu Z, Brignardello J, Kao D, Holmes E, Li JV, Clarke TB, Thursz MR, Marchesi JR. 2018. Inhibiting growth of clostridioides difficile by restoring valerate, produced by the intestinal microbiota. Gastroenterology 155:1495–1507.e15. doi:10.1053/j.gastro.2018.07.014.

17. Ghimire S, Roy C, Wongkuna S, Antony L, Maji A, Keena MC, Foley A, Scaria J. 2020. Identification of clostridioides difficile-inhibiting gut commensals using culturomics, phenotyping, and combinatorial community assembly. mSystems 5. doi:10.1128/msystems.00620-19.

18. Auchtung JM, Preisner EC, Collins J, Lerma AI, Britton RA. 2020. Identification of simplified microbial communities that inhibit clostridioides difficile infection through dilution/extinction. mSphere 5. doi:10.1128/msphere.00387-20.

19. Schubert AM, Sinani H, Schloss PD. 2015. Antibiotic-induced alterations of the murine gut microbiota and subsequent effects on colonization resistance against clostridium difficile. mBio 6. doi:10.1128/mbio.00974-15.

20. Jenior ML, Leslie JL, Young VB, Schloss PD. 2018. Clostridium difficile alters the structure and metabolism of distinct cecal microbiomes during initial infection to promote sustained colonization. mSphere 3. doi:10.1128/msphere.00261-18.

21. Kurtz ZD, Müller CL, Miraldi ER, Littman DR, Blaser MJ, Bonneau RA. 2015. Sparse and compositionally robust inference of microbial ecological networks. PLOS Computational Biology 11:e1004226. doi:10.1371/journal.pcbi.1004226.

22. Reeves AE, Theriot CM, Bergin IL, Huffnagle GB, Schloss PD, Young VB. 2011. The interplay between microbiome dynamics and pathogen dynamics in a murine model ofClostridium difficileInfection. Gut Microbes 2:145–158. doi:10.4161/gmic.2.3.16333.

23. Reeves AE, Koenigsknecht MJ, Bergin IL, Young VB. 2012. Suppression of clostridium difficile in the gastrointestinal tracts of germfree mice inoculated with a murine isolate from the family lachnospiraceae. Infection and Immunity 80:3786–3794. doi:10.1128/iai.00647-12.

24. Tomkovich S, Stough JMA, Bishop L, Schloss PD. 2020. The initial gut microbiota and response to antibiotic perturbation influence clostridioides difficile clearance in mice. mSphere 5. doi:10.1128/msphere.00869-20.

25. Peterfreund GL, Vandivier LE, Sinha R, Marozsan AJ, Olson WC, Zhu J, Bushman FD. 2012. Succession in the gut microbiome following antibiotic and antibody therapies for clostridium difficile. PLoS ONE 7:e46966. doi:10.1371/journal.pone.0046966.

26. Fukuyama J, Rumker L, Sankaran K, Jeganathan P, Dethlefsen L, Relman DA, Holmes SP. 2017. Multidomain analyses of a longitudinal human microbiome intestinal cleanout perturbation experiment. PLOS Computational Biology 13:e1005706. doi:10.1371/journal.pcbi.1005706.

27. Suez J, Zmora N, Zilberman-Schapira G, Mor U, Dori-Bachash M, Bashiardes S, Zur M, Regev-Lehavi D, Brik RB-Z, Federici S, Horn M, Cohen Y, Moor AE, Zeevi D, Korem T, Kotler E, Harmelin A, Itzkovitz S, Maharshak N, Shibolet O, Pevsner-Fischer M, Shapiro H, Sharon I, Halpern Z, Segal E, Elinav E. 2018. Post-antibiotic gut mucosal microbiome reconstitution is impaired by probiotics and improved by autologous FMT. Cell 174:1406–1423.e16. doi:10.1016/j.cell.2018.08.047.

28. Ianiro G, Maida M, Burisch J, Simonelli C, Hold G, Ventimiglia M, Gasbarrini A, Cammarota G. 2018. Efficacy of different faecal microbiota transplantation protocols for clostridium difficile infection: A systematic review and meta-analysis. United European Gastroenterology Journal 6:1232–1244. doi:10.1177/2050640618780762.

29. Allegretti JR, Mehta SR, Kassam Z, Kelly CR, Kao D, Xu H, Fischer M. 2020. Risk factors that predict the failure of multiple fecal microbiota transplantations for clostridioides difficile infection. Digestive Diseases and Sciences. doi:10.1007/s10620-020-06198-2.

30. Garza-González E, Mendoza-Olazarán S, Morfin-Otero R, Ramírez-Fontes A, Rodríguez-Zulueta P, Flores-Treviño S, Bocanegra-Ibarias P, Maldonado-Garza H, Camacho-Ortiz A. 2019. Intestinal microbiome changes in fecal microbiota transplant (FMT) vs. FMT enriched with lactobacillus in the treatment of recurrent clostridioides difficile infection. Canadian Journal of Gastroenterology and Hepatology 2019:1–7. doi:10.1155/2019/4549298.

31. Winter SE, Lopez CA, Bäumler AJ. 2013. The dynamics of gut-associated microbial communities during inflammation. EMBO reports 14:319–327. doi:10.1038/embor.2013.27.

32. Rivera-Chávez F, Lopez CA, Bäumler AJ. 2017. Oxygen as a driver of gut dysbiosis. Free Radical Biology and Medicine 105:93–101. doi:10.1016/j.freeradbiomed.2016.09.022.

33. Carlson PE, Walk ST, Bourgis AET, Liu MW, Kopliku F, Lo E, Young VB, Aronoff DM, Hanna PC. 2013. The relationship between phenotype, ribotype, and clinical disease in human clostridium difficile isolates. Anaerobe 24:109–116. doi:10.1016/j.anaerobe.2013.04.003.

34. Thanissery R, Winston JA, Theriot CM. 2017. Inhibition of spore germination, growth, and toxin activity of clinically relevant c. difficile strains by gut microbiota derived secondary bile acids. Anaerobe 45:86–100. doi:10.1016/j.anaerobe.2017.03.004.

35. Theriot CM, Koumpouras CC, Carlson PE, Bergin II, Aronoff DM, Young VB. 2011. Cefoperazone-treated mice as an experimental platform to assess differential virulence ofClostridium difficilestrains. Gut Microbes 2:326–334. doi:10.4161/gmic.19142.

36. Rao K, Micic D, Natarajan M, Winters S, Kiel MJ, Walk ST, Santhosh K, Mogle JA, Galecki AT, LeBar W, Higgins PDR, Young VB, Aronoff DM. 2015. Clostridium difficile Ribotype 027: Relationship to age, detectability of toxins a or b in stool with rapid testing, severe infection, and mortality. Clinical Infectious Diseases 61:233–241. doi:10.1093/cid/civ254.

37. Sorg JA, Dineen SS. 2009. Laboratory maintenance ofClostridium difficile. Current Protocols in Microbiology 12. doi:10.1002/9780471729259.mc09a01s12.

38. Winston JA, Thanissery R, Montgomery SA, Theriot CM. 2016. Cefoperazone-treated mouse model of clinically-relevant Δclostridium difficile strain r20291. Journal of Visualized Experiments. doi:10.3791/54850.

39. Kozich JJ, Westcott SL, Baxter NT, Highlander SK, Schloss PD. 2013. Development of a dual-index sequencing strategy and curation pipeline for analyzing amplicon sequence data on the MiSeq illumina sequencing platform. Applied and Environmental Microbiology 79:5112–5120. doi:10.1128/aem.01043-13.

40. Schloss PD, Westcott SL, Ryabin T, Hall JR, Hartmann M, Hollister EB, Lesniewski RA, Oakley BB, Parks DH, Robinson CJ, Sahl JW, Stres B, Thallinger GG, Horn DJV, Weber CF. 2009. Introducing mothur: Open-source, platform-independent, community-supported software for describing and comparing microbial communities. Applied and Environmental Microbiology 75:7537–7541. doi:10.1128/aem.01541-09.

41. Wang Q, Garrity GM, Tiedje JM, Cole JR. 2007. Naïve bayesian classifier for rapid assignment of rRNA sequences into the new bacterial taxonomy. Applied and Environmental Microbiology 73:5261–5267. doi:10.1128/aem.00062-07.

42. Yue JC, Clayton MK. 2005. A similarity measure based on species proportions. Communications in Statistics - Theory and Methods 34:2123–2131. doi:10.1080/sta-200066418.

43. Benjamini Y, Hochberg Y. 1995. Controlling the false discovery rate: A practical and powerful approach to multiple testing. Journal of the Royal Statistical Society: Series B (Methodological) 57:289–300. doi:10.1111/j.2517-6161.1995.tb02031.x.

44. Topçuoǧlu BD, Lesniak NA, Ruffin MT, Wiens J, Schloss PD. 2020. A framework for effective application of machine learning to microbiome-based classification problems. mBio 11. doi:10.1128/mbio.00434-20.

